# The Transcription Factor Foxi1 Promotes Expression of V-ATPase and Gpr116 in M-1 cells

**DOI:** 10.1101/2022.10.31.514579

**Authors:** Mackenzie Kui, Jennifer L. Pluznick, Nathan A. Zaidman

**Affiliations:** Department of Physiology, Johns Hopkins University School of Medicine; Department of Biochemistry and Molecular Biology, University of New Mexico School of Medicine

**Keywords:** Foxi1, V-ATPase, Gpr116, intercalated cells, collecting duct

## Abstract

The diverse functions of each nephron segment rely on the coordinated action of specialized cell populations that are uniquely defined by their transcriptional profile. In the collecting duct, there are two critical and distinct cell populations: principal cells and intercalated cells. Principal cells play key roles in the regulation of water, Na^+^, and K^+^, while intercalated cells are best known for their role in acid-base homeostasis. Currently, there are no *in vitro* systems that recapitulate the heterogeneity of the collecting ducts, which limits high-throughput and replicate investigations of genetic and physiological phenomena. Here, we have demonstrated that the transcription factor Foxi1 is sufficient to alter the transcriptional identity of M-1 cells, a murine cortical collecting duct cell line. Specifically, overexpression of *Foxi1* induces the expression of intercalated cell transcripts including *Gpr116, Atp6v1b1, Atp6v1g3, Atp6v0d2, Slc4a9*, and *Slc26a4*. These data indicate that overexpression of *Foxi1* differentiates M-1 cells towards a B-type intercalated cell phenotype and may provide a novel *in vitro* tool to study transcriptional regulation and physiological function of the renal collecting duct.

## Introduction

The renal collecting duct is a heterogenous epithelium comprised of principal cells (PCs) and intercalated cells (ICs). PCs play a critical role in the maintenance of blood volume and are sensitive to both vasopressin and aldosterone, which drive apical expression of aquaporin-2 (AQP2) water channels and epithelial sodium channels (ENaC), respectively (1). Intercalated cells are characterized by plasma membrane expression of the vacuolar-type H^+^-ATPase (V-ATPase); A-type ICs (AICs) express V-ATPase on the luminal membrane, and B-type ICs (BICs) express V-ATPase on the serosal membrane (2, 3). Due to their reliance on V-ATPase for transport processes, ICs are considered important renal regulators of pH homeostasis.

Unlike other epithelial tissues (such as the lung (4, 5) and intestine (6, 7)) there are limited *in vitro* systems that recapitulate the physiological function of the nephron, let alone the patterned collecting ducts (though a recent report demonstrated directed differentiation of human pluripotent stem cells into collecting duct epithelial cells (8)). To date, significant advances in our understanding of transport phenomena in the collecting ducts has relied on low-throughput methods such as perfused (9, 10) or split-open single tubules (11, 12), as well as transgenic animal models. Immortalized cell lines, such as mpkCCD cells, have been effective tools in understanding AQP2 regulation in PCs (13, 14), as well as amiloride-sensitive ENaC current (15, 16). However, these cells vary by clonal population and are largely representative of PCs, not ICs. Therefore, an *in vitro* system that replicates the patterned cell types of the collecting duct could become an important tool for the investigation of both PC and IC physiology.

The cells of the collecting duct are derived developmentally from the ureteric bud, which mature in a Notch-dependent manner into the “salt and pepper” distribution of minority ICs among majority PCs (17, 18). Genetic deletion of the transcription factor Foxi1 leads to the absence of ICs, and Foxi1 deficient mice develop distal renal tubular acidosis (19, 20). Foxi1 mutation is similarly associated with acidosis and deafness in humans (21). Indeed, Foxi1 is a transcriptional regulator of IC-specific expression of V-ATPase subunits (22, 23) as well as BIC specific expression of AE4 (24), a sodium-dependent anion exchanger, emphasizing the critical role of Foxi1 in IC specification. Additionally, a recent study identified Foxi1 as a critical regulator of airway CFTR-rich ionocytes (25). Notably, these pulmonary ionocytes express an adhesion-class G protein-coupled receptor (Gpr116/ADGRF5) that we previously revealed to be a critical regulator of V-ATPase in AICs (26), suggesting that Foxi1 initiates a transcriptional program that includes Gpr116. Here, we test the hypothesis that heterologous expression of *Foxi1* in an immortalized murine cortical collecting duct cell line (M-1 (27)) is sufficient to generate IC-like cells *in vitro*.

## Methods

### Cell culture

M-1 cells (ATCC, VA) were grown in a 1:1 mixture of Dulbecco’s modified Eagle’s medium and Ham’s F12 medium with 2.5 mM L-glutamine adjusted to contain 15 mM HEPES, 0.5 mM sodium pyruvate and 1.2 g/L sodium bicarbonate supplemented with 0.005 mM dexamethasone, 10% fetal bovine serum, 1% P/S and 5 μg/mL plasmocin (Invivogen, CA). M-1 cells were grown on coverslips (for RNAScope) or Transwell 0.4 μm pore polyester membranes (Corning, AZ). HEK29T, MCF7, IMCD3, MDCK, and MDCK-C11 cells were grown in MEM supplemented with 10% FBS, 1% P/S and 1% L-glutamine. Cells were maintained in a 37°C humidified incubator with 5% CO_2_. Once cultures reached 60% confluence, cells were reverse transfected with Lipofectamine 2000 (Thermo Fisher Scientific, MA) at a plasmid mass to transfection reagent volume ratio of 2:1 and grown for at least two days prior to experimentation.

### Cloning

*Foxi1* (NM_023907.4) cDNA was cloned from whole mouse kidney into a pME18S expression vector (pFoxi1) using the forward primer 5’ TAATAG<u>gaattc</u>ATGAGCTCCTTCGAC 3’ (EcoRI) and reverse primer 5’ TAATTT<u>ctcgag</u>CTAGACTTCAGTGCC 3’ (XhoI) and validated by Sanger sequencing. *Foxi1* was then subcloned into the pLVX-EF1alpha-IRES-mCherry vector at EcoRI and BamHI restriction sites using forward primer 5’ TAATAG<u>gaattc</u>ATGAGCTCCTTCGAC 3’ and reverse primer 5’ TAATTT<u>ggatcc</u>CTAGACTTCAGTGCC 3’ to make pFoxi1-IRES-mCherry.

### Conventional and Quantitative RT-PCR

RNA was isolated from cells and mouse kidney using the RNeasy Mini Kit (Qiagen, DE). cDNA was generated using the QuantiTect reverse transcription kit with gDNA Wipeout (Qiagen, DE). Conventional PCR amplification of *Gpr116* was achieved using the forward primer 5’ TCCAATTCGAGGGACCGAAG 3’ and reverse primer 5’ GTAGTTCACAACCACGCTGC 3’. TaqMan real-time PCR assays were performed in triplicate according to the manufacturer’s standard protocol (Thermo). 10 ng cDNA was PCR-amplified using an Applied Biosystems QuantStudio 6 PCR system (Thermo). Amplification data was analyzed on Microsoft Excel and plotted with GraphPad Prism version 9.1.2. All qPCR data is presented as relative expression per *Gapdh*. TaqMan assays are listed in Table 1.

**Table 1:** TaqMan Assays and RNAscope Probes.

| Target | TaqMan Assay ID |
| --- | --- |
| AQP2 | Mm00437575_m1 |
| AQP6 | Mm00558232_m1 |
| ATP6V0A4 | Mm00459882_m1 |
| ATP6V0D2 | Mm01222963_m1 |
| ATP6V1B1 | Mm00460309_m1 |
| ATP6V1G3 | Mm00616840_m1 |
| AVPR2 | Mm01193534_g1 |
| CAR2 | Mm00501576_m1 |
| CFTR | Mm00445197_m1 |
| CLCN5 | Mm00443851_m1 |
| DMRT2 | Mm00659912_m1 |
| GPR116 | Mm01269030_m1 |
| SCNN1A | Mm00803386_m1 |
| SLC26A4 | Mm01258316_m1 |
| SLC4A1 | Mm01245920_g1 |
| SLC4A9 | Mm01130729_m1 |
| GAPDH | Mm99999915_g1 |
| Target | RNAscope Probe |
| FOXI1 | 483021 |
| GPR116 | 318021-C2 |

### Immunofluorescent Microscopy and RNAscope In Situ Hybridization

Mice harboring an EGFP gene driven by the ATP6V1B1 promoter (kind gift from Dennis Brown, MGH (28)) were perfusion-fixed by intracardial transfusion of approximately 20mL of PBS followed by 20mL of 4% PFA under deep anesthesia. Kidneys were harvested and drop-fixed in 4% PFA overnight at 4°C. The kidneys were then dehydrated in a 30% sucrose solution overnight, embedded into Tissue-Tek O.C.T. compound, and flash frozen with liquid nitrogen. Frozen kidneys were sectioned at 8 μm on a cryostat and mounted on slides.

M-1 cells were grown on Transwell membranes (Immunofluorescence, IF) or Nunc Lab-Tek chambered coverglass (RNAscope, Thermo) until 60% confluent, transfected with pFoxi1 or pFoxi1-IRES-mCherry then fixed with 4% PFA after four days (IF) or two days (RNAscope) of culture. For IF, cells were permeabilized with 0.3% Triton-X for 10 minutes then blocked with 3% BSA for 60 minutes. Cells were treated with primary antibodies (Foxi1: ab20454, Abcam; V-ATPaseβ1/2: sc-55544, Santa Cruz Biotechnology) diluted 1:100 in blocking buffer overnight at 4°C. Cells were washed in PBS and exposed to secondary antibodies diluted 1:500 in PBS for 45 minutes at room temperature, then counterstained with DAPI. Membranes were excised from the Transwells and mounted on glass slides for imaging.

RNAscope *in situ* hybridization was performed using the RNAscope Fluorescent Multiplex Assay kit per manufacturer’s instructions (Biotechne, MN). RNAscope hybridization probes specific to murine *Foxi1* and *Gpr116* are listed in Table 1. To improve detection of EGFP signal after RNAscope, a primary antibody targeting GFP was added (diluted 1:100 in blocking buffer, A-6455, Thermo), followed by addition of a fluorescent Alexa Fluor secondary antibody. Fluorescent images were captured on a Zeiss LSM880-Airyscan FAST Super-Resolution laser scanning confocal microscope and processed in Zeiss Zen software at the Johns Hopkins University School of Medicine Microscope Core Facility.

### Statistics

All values are given as mean±SEM. Student’s t-test was used for statistical analysis. A p-value of less than 0.05 was considered to be significant.

## Results

### Foxi1 and Gpr116 are expressed in the renal collecting duct

To verify that Foxi1 and Gpr116 are co-expressed in renal intercalated cells, we used transgenic mice expressing EGFP driven by the *Atp6v1b1* promoter to identify V-ATPase expressing intercalated cells in the collecting ducts (Figure 1). RNAscope *in situ* hybridization confirmed expression of *Foxi1* and *Gpr116* in EGFP-expressing cells, though a subset of cells expresses *Foxi1* without *Gpr116*.

**Figure 1:**
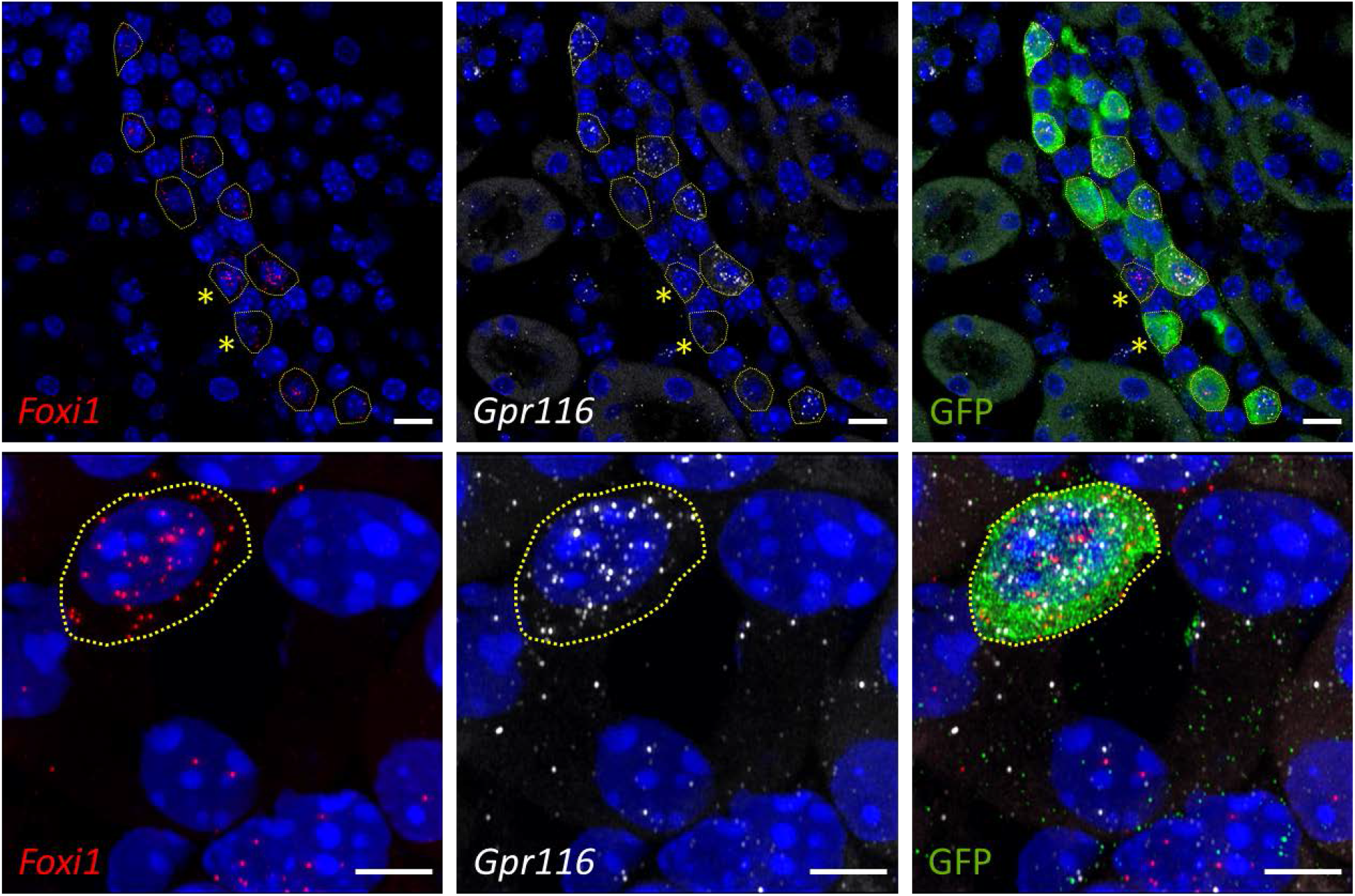
Co-expression of *Foxi1* and *Gpr116* in murine collecting ducts. Detection of *Foxi1* (red) and *Gpr116* (white) transcripts by RNAscope hybridization demonstrates co-localization in intercalated cells (identified by EGFP expression in ATP6V1B1-EGFP mice). *Gpr116* is not detected in some Foxi1^+^ cells (top panels, asterisk). Scale bars = 10 μm (top), 5 μm (bottom).

### M-1 cells transfected with pFoxi1 induce expression of Gpr116

Next, we overexpressed murine *Foxi1* in immortalized cell lines (Figure 2A). RT-PCR revealed an induction of *Gpr116* transcript in M-1 cells, but not in HEK293T, IMCD3, MCF7, MDCK or MDCK-C11 cells (Figure 2A/B). These results were validated by qPCR in M-1 cells (Figure 2C). M-1 cells do not express endogenous *Foxi1*, but M-1 cells transfected with pFoxi1 express *Gpr116* as detected by RNAscope (Figure 2D).

**Figure 2:**
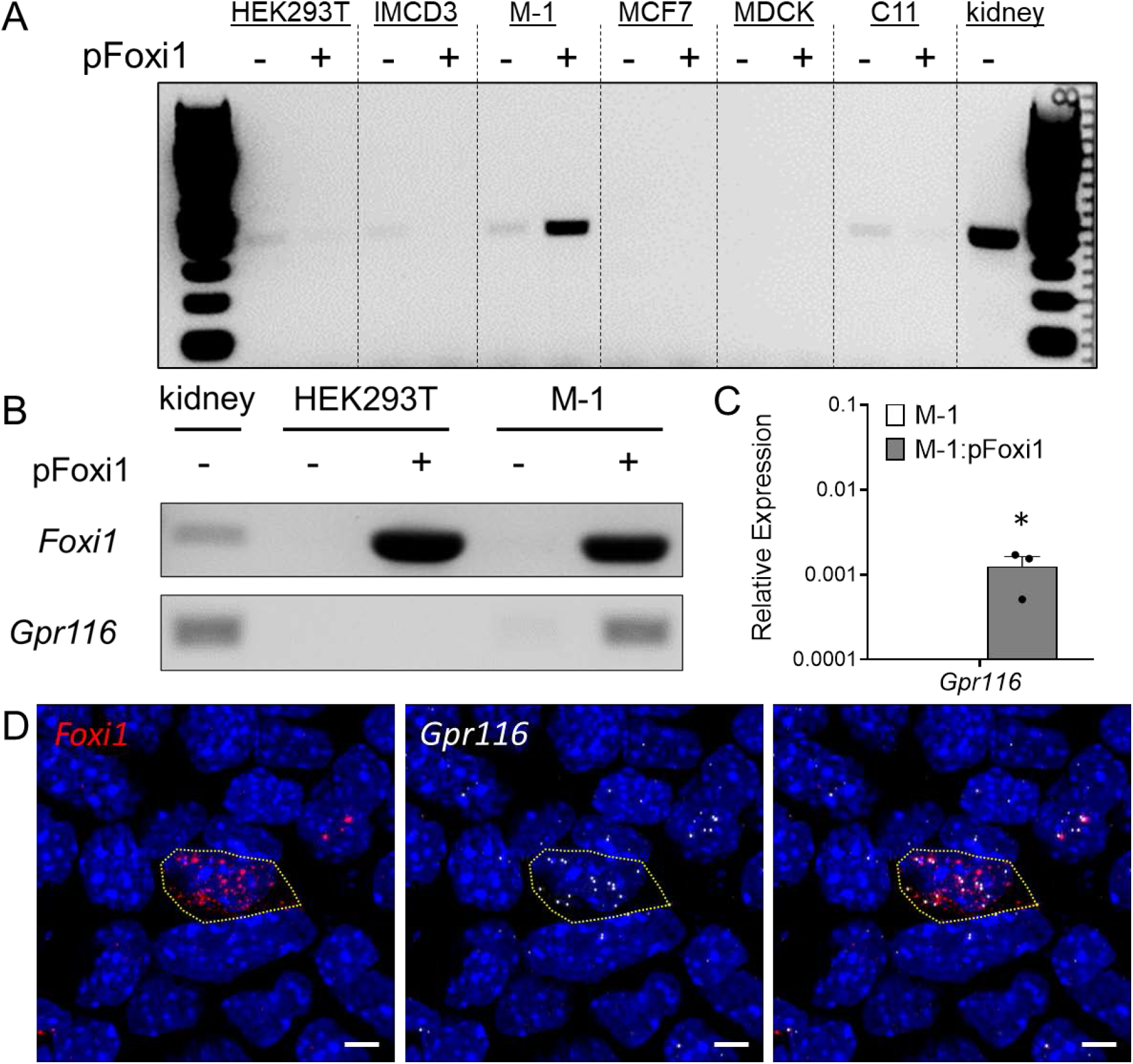
Induction of Gpr116 transcription in M-1 cells transfected with Foxi1. (A) Heterologous expression of *Foxi1* induces *Gpr116* transcription in M-1 cells but not in other immortalized cell lines, as revealed by RT-PCR. Endogenous expression of *Gpr116* in murine kidney is shown as a positive control. (B) Repeat measurement of *Gpr116* expression in M-1 cells transfected with pFoxi1. Endogenous expression of *Foxi1* and *Gpr116* in murine kidney is shown as a positive control (C) *Foxi1-*induced *Gpr116* transcription in M-1 cells as measured by Taqman qPCR. Relative expression is per *Gapdh*. Cycle threshold (C_T_) of 35≈Relative Expression 0.0001. N=3. Bars are mean±SEM. \**P*<0.05 vs. M-1. (D) Co-localization of *Foxi1* and *Gpr116* transcripts in an M-1 cell transfected with pFOXI1. Scale bar = 5μm.

### Foxi1 overexpression in M-1 cells induces V-ATPase subunit transcription

*Foxi1* is a transcriptional regulator of V-ATPase proton pump subunits and a necessary transcription factor for intercalated cell specification. Therefore, we investigated if transfection with *Foxi1* caused a broader remodeling of the transcriptional landscape in M-1 cells. Indeed, we observed induction of the *Atp6v1b1, Atp6v1g3*, and *Atp6v0d2* V-ATPase subunits in M-1 cells (Figure 3A). Notably, M-1 cells have no endogenous expression of these V-ATPase subunits. Additionally, M-1 cells express endogenous *Atp6v0a4* and this transcript was not affected by *Foxi1* transfection.

**Figure 3:**
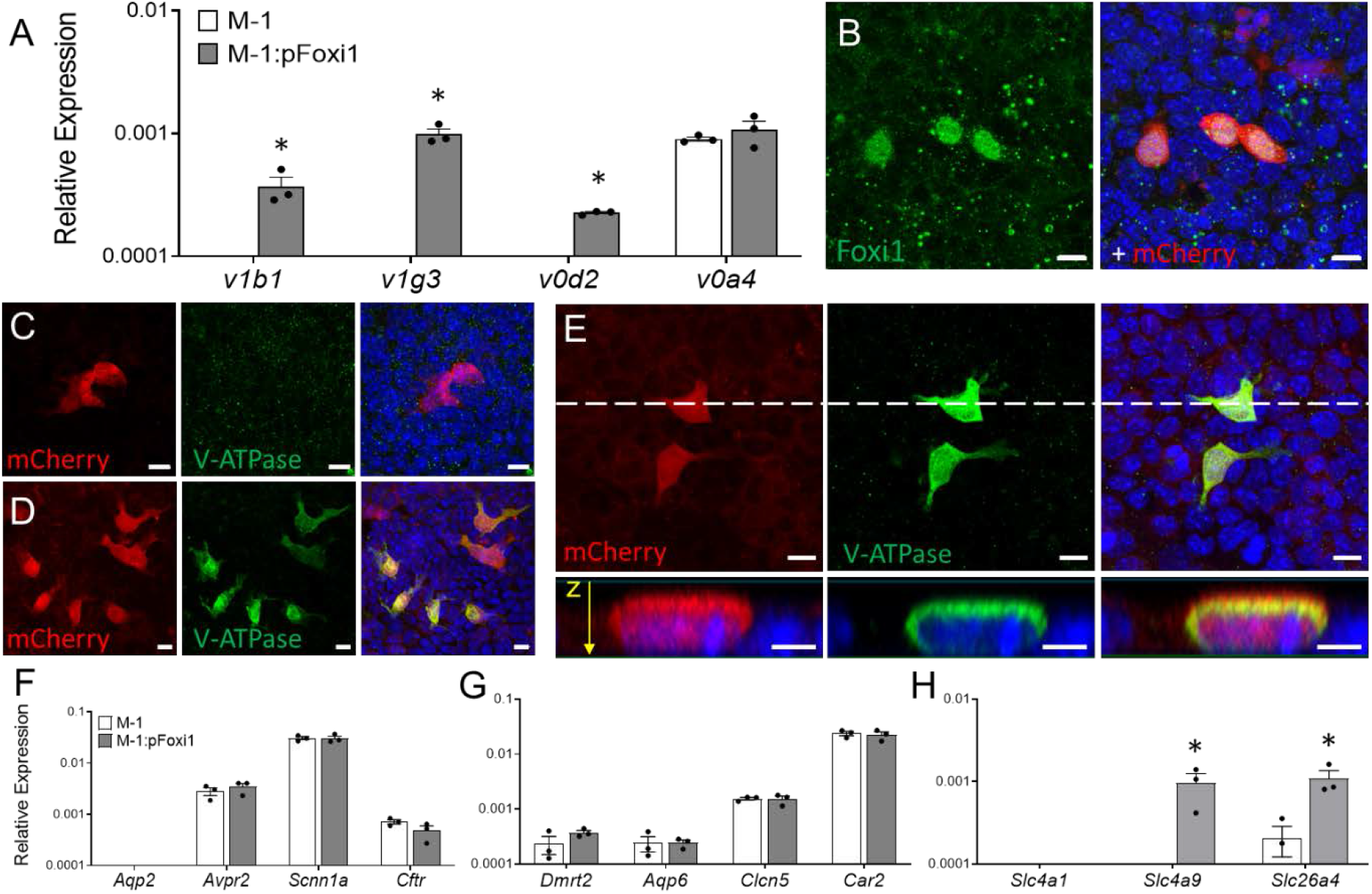
Transfection with pFoxi1 induces V-ATPase subunit transcription in M-1 cells. (A) Heterologous expression of *Foxi1* induces transcription of V-ATPase subunits *Atp6v1b1, Atp6v1g3*, and *Atp6v0d2* as measured by Taqman qPCR. (B) M-1 cells transfected with pFoxi1-IRES-mCherry express *Foxi1* as detected by immunofluorescent microscopy (IF). (C) M-1 cells transfected with control IRES-mCherry vector do not express V-ATPase. (D) M-1 cells transfected with pFoxi1-IRES-mCherry express V-ATPase β1/2 subunit as detected by IF. (E) V-ATPase localizes at or near the plasma membrane in transfected M-1 cells but is not restricted to the apical or basolateral side. Dashed line is orthogonal plane shown at bottom. Z-plane is labeled from apical to basolateral; scale bar = 10μm. (F) Transcripts associated with principal cells are unaffected by pFoxi1. (G) Many transcripts associated with intercalated cells are unaffected by pFoxi1. (H) pFoxi1 induces *Slc4a9* (AE4) and *Slc26a4* (Pendrin) transcription in M-1 cells as measured by Taqman qPCR. Relative expression is per *Gapdh*. Cycle threshold (C_T_) of 35≈Relative Expression 0.0001. N=3. Bars are mean±SEM. \**P*<0.05 vs. M-1. All scale bars = 5μm unless noted (orthogonal images in E).

Next, using a heterologous expression vector containing *Foxi1* and mCherry separated by an internal ribosome entry site (pFoxi1-IRES-mCherry, Figure 3B), we confirmed expression of V-ATPase protein in transfected M-1 cells by immunofluorescent microscopy (Figures 3C/D/E). V-ATPase appears to localize at or near the plasma membrane in M-1 cells grown on Transwell filter membranes. V-ATPase localization was not polarized towards the apical or basolateral membranes (Figure 3E).

### Foxi1 induced transcription of Ae4 and Pendrin

Several other genes associated with PCs (*Aqp2, Avpr2, Scnn1a, Cftr*) and ICs (*Dmrt2, Aqp6, Clcn5, Car2*) were unchanged by qPCR in response to *Foxi1* overexpression (Figures 3F/G). However, M-1 cells transfected by pFoxi1 demonstrated transcriptional upregulation of *Slc4a9* (Ae4) and *Slc26a4* (Pendrin), both specific transcripts of BICs (Figure 3H). In contrast, the AIC-specific transcript *Slc4a1* (Ae1) was not detected in transfected cells.

## Discussion

We hypothesized that heterologous expression of the transcription factor *Foxi1* was sufficient to generate IC-like cells *in vitro*. Here we present an *in vitro* system that transforms M-1 collecting duct cells into heterogenous monolayers harboring cells that express *Gpr116, Atp6v1b1* and *Slc4a9*. Though considerable work remains to functionally characterize our system, the generation of cells expressing IC-specific transcripts represents a significant development for high-throughput analysis of collecting duct physiology. Of note, pFoxi1 does not induce transcription of the AIC-specific *Slc4a1* (Ae1) in M-1 cells. This implies that our *in vitro* system is a model of BIC-like cells, characterized by *Slc4a9* expression. Though Gpr116 is restricted to AICs *in vivo*, it may be that Gpr116 expression is initiated by Foxi1 during IC specification, with further layers of regulation contributing to Gpr116 expression in A versus BICs. Our results demonstrate that there are additional factors required to generate AIC-specific transcripts like *Slc4a1*. Whether Foxi1 is necessary and permissible for *Slc4a1* expression is the subject of ongoing investigation.

Interestingly, heterologous expression of *Foxi1* does not induce transcription of Gpr116 or V-ATPase subunits in all cell lines. We tested several immortalized cell lines including IMCD3, MCF7, MDCK, MDCK C11, and HEK293T, but only M-1 demonstrated a transcriptional response to pFoxi1. M-1 cells are reported to have both PC and IC-like characteristics (27). Indeed, transcriptionally, M-1 cells express PC-specific genes such as *Avpr2* and *Scnn1a*, as well as *Dmrt2, Car2* and *Aqp6* (IC-specific markers). Our finding that pFoxi1 induces *Atp6v1b1* transcription in M-1 cells suggests the epigenetic landscape of M-1 cells is permissive to transcriptional regulation by Foxi1.

Though M-1 cells are clearly responsive to Foxi1 transcriptional regulation, we do not yet have a full understanding of the extent of IC-specification in cultured cells. While we can detect robust upregulation of *Foxi1* target genes (*Atp6v1b1, Gpr116, Slc4a9*) by qPCR, this transcriptional activity presumably occurs only in the subset of M-1 cells that are transfected by pFoxi1. And, since the majority of M-1 cells in a culture dish are not transfected and express PC-specific genes such as *Avpr2*, we are unable to determine if pFoxi1 causes the downregulation of PC genes in transfected M-1 cells. Therefore, our data does not eliminate the possibility that pFoxi1 transfected M-1 cells downregulate PC-specific genes in favor of BIC-specific Foxi1 target genes.

In addition to inducing V-ATPase transcription, M-1 cells transfected with pFoxi1 significantly upregulate Slc4a9 (Ae4) and Slc26a4 (Pendrin). The expression of Slc4a9 is in agreement with previous studies showing that Slc4a9 is targeted by Foxi1 (24). The generation of cells expressing Ae4, a poorly understood Cl^-^/HCO_3_^-^ exchanger, represents a potentially significant advancement for *in vitro* investigations of Ae4 and BIC function. Future experiments that assess the functional attributes of these cells will be critical to determining the usefulness of this cell model, which we hope will become a useful tool for investigations of IC differentiation, function and regulation.

## Acknowledgements

The authors acknowledge the members of the Pluznick lab for constructive discussion and support, as well as the faculty, trainees and staff of the Department of Physiology and Johns Hopkins School of Medicine Microscopy Facility. We also acknowledge Dennis Brown (MGH) for kindly providing EGFP-reporter mice.

## Author Contributions

M.K. and N.A.Z. developed the concept and designed the research; M.K. and N.A.Z. performed the experiments; M.K. and N.A.Z. analyzed the data; M.K., J.L.P. and N.A.Z. interpreted the results of the experiments; M.K. and N.A.Z. prepared the figures; M.K. and N.A.Z. drafted the manuscript; M.K., J.L.P. and N.A.Z. edited and revised the manuscript; M.K., J.L.P. and N.A.Z. approved the final version of the manuscript.

